# Disease-attenuated pneumococcal biosynthesis gene mutants invade the mucosal epithelium and induce innate immunity

**DOI:** 10.1101/2023.06.15.545009

**Authors:** Caroline M. Weight, Gabriele Pollara, Modupeh Betts, Roberta Ragazzini, Elisa Ramos-Sevillano, Jesús Reiné, Matthew Whelan, José Afonso Guerra-Assunção, Michael Connor, Paola Bonfanti, Clare Jolly, Mahdad Noursadeghi, Daniela M. Ferreira, Jeremy S. Brown, Robert S. Heyderman

**Author notes:** Corresponding Authors: Dr Caroline M. Weight and Professor Robert S. Heyderman, and. Current address Biomedical and Life Sciences, Faculty of Health and Medicine, Lancaster University, UK.

## Abstract

Nasopharyngeal colonisation by *Streptococcus pneumoniae* is characterised by bacterial adherence to epithelial cells, microinvasion and innate immune activation. Previously, we have shown two serotype 6B *S. pneumoniae* mutant strains affecting bacterial metabolism (*ΔproABC/pia* and *Δfhs/pia*) colonise humans and mice, but in a murine disease model do not cause invasive infection. Here, we explore whether *S. pneumoniae* epithelial microinvasion and the induction of innate immune responses persist despite disease attenuation.

We show that under serum stress, these biosynthesis gene mutations had a broad but different impact on pneumococcal virulence gene expression, oxidative stress regulation, and purine and carbohydrate metabolism genes. However, although these mutations did not attenuate microinvasion in human challenge and epithelial models, there was less transmigration of Detroit 562 nasopharyngeal epithelial cells by the mutants compared to WT. Cellular reorganisation of primary human airway epithelium varied considerably between strains. Compared to WT, infection of Detroit 562 epithelial cells by the *Δfhs/piaA* strain, but not the *ΔproABC/piaA* strain was less pro-inflammatory, induced less caspase 8 production, and were associated with increased pneumococcal hydrogen peroxide and reduced pneumolysin secretion.

These findings suggest that the observed differences in microinvasion and the epithelial response were driven by the differential expression of multiple bacterial virulence and metabolic pathways, rather than single genes or pathways of genes. These data highlight the complex impact of single gene mutations on bacterial virulence and suggest that the virulence determinants of pneumococcal epithelial colonisation, microinvasion and innate immunity are not necessarily directly linked to disease.

**Author Summary:** *Streptococcus pneumoniae* (the pneumococcus) commonly colonises the back of the human nose, and is a leading cause of pneumonia, meningitis, and sepsis. During colonisation, the pneumococcus adheres to the cells in the nose, invades these cells (so-called microinvasion), and activates them. Colonisation is a pre-requisite for disease, however, since disease is largely a dead end for *S. pneumoniae*, it remains unclear whether these processes are directly linked to disease progression. We have previously shown that if we introduce gene mutations into *S. pneumoniae* that affect key metabolic pathways, these bacteria retain their ability to colonize human and animal models without causing disease. We now show that these mutants retain their ability to microinvade epithelial cells in human and mouse models, and some may still cause inflammation, but are less able to pass through the epithelial barrier. However, although the attenuation of disease may be explained by the broad-ranging impact of these mutations on pneumococcal virulence, oxidative stress, and metabolism, they are not driven by a single determinant. Our findings suggest that pneumococcal microinvasion and immune activation are not necessarily pre-cursors to disease progression. This supports the idea that *S. pneumoniae* adapts and evolves to promote colonisation and ultimately transmission rather than cause disease.

**Graphical Abstract:** *S. pneumoniae* colonisation is characterised by mucus association, epithelial adherence, microcolony formation and microinvasion – where the pneumococcus invades the epithelial barrier without causing disease. Although mutations in *S. pneumoniae* biosynthesis genes (*ΔproABC* and *Δfhs*) attenuate disease in a murine model, they do not attenuate microinvasion in either experimental human pneumococcal challenge (EHPC), *ex vivo* or *in vitro* epithelial cells. Transmigration of the epithelial barrier is attenuated. These mutations show strain-dependent effects on both the epithelial and bacterial responses to infection. Factors such as epithelial cellular reorganisation, inflammation and caspase 8 activity alongside pneumococcal metabolic adaptation, virulence factor expression and response to stress are important components of these processes.

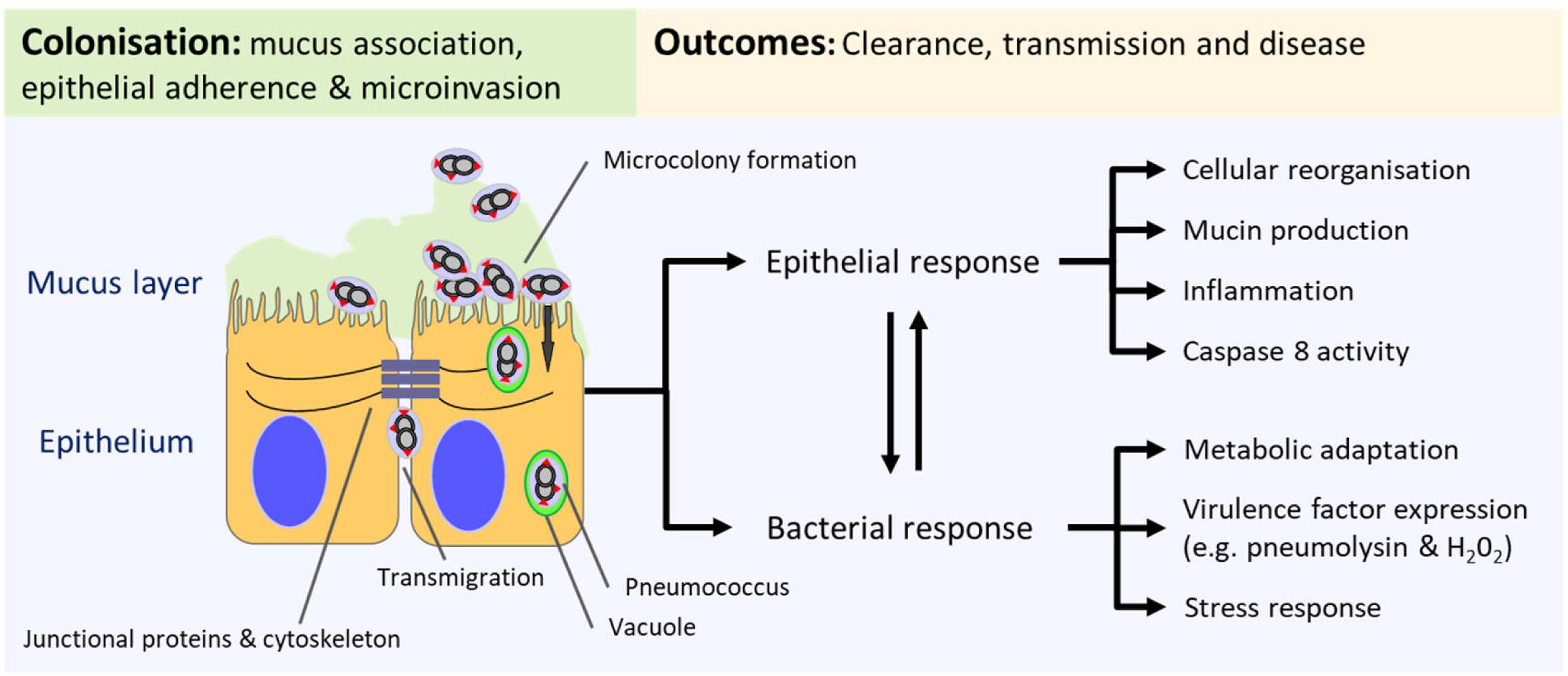

## INTRODUCTION

*Streptococcus pneumoniae* is a common commensal of the URT, yet translocation of the pneumococcus to the lungs, blood or the brain leads to life-threatening disease [1]. Among children under five, *S. pneumoniae* is the leading cause of pneumonia-related deaths globally [2].

We have previously shown that *S. pneumoniae* colonisation is characterised by epithelial surface microcolony formation and invasion, termed microinvasion [3]. Pneumococcal colonization elicits an innate immune response that may facilitate clearance while also promoting bacterial transmission [3, 4]. Beyond the capsule, numerous virulence factors have evolved to optimise colonization and transmission, rather to cause invasive disease. We therefore hypothesize that epithelial microinvasion and the innate immune response to colonization are not necessarily linked to disease causation.

To test this, we have used two *S. pneumoniae* serotype 6B biosynthesis gene mutants (Δ*proABC* and Δ*fhs*), which were designed to induce protective immunity through nasopharyngeal colonization without causing disease [5]. The *proABC* operon encodes enzymes for proline biosynthesis, while *fhs* encodes a tetrahydrofolate reductase involved in one-carbon metabolism, both essential for pneumococcal growth and virulence [6–8]. In murine models, these mutants colonize the nasopharynx but fail to cause invasive disease [5, 8]. In EHPC, Δ*proABC* and Δ*fhs* mutants carrying an additional *piaA* deletion (disrupting a key iron uptake transporter, introduced to prevent reversion) retained the ability to colonise and elicit IgG-mediated immunity [9, 10], [10]. These isogenic mutants thus provide a controlled system to investigate epithelial microinvasion and innate immune activation independently of capsular serotype.

Here, we show that although the Δ*proABC/piaA* and Δ*fhs/piaA* mutations attenuate disease through broad-ranging impacts on pneumococcal virulence, oxidative stress, and metabolism, and reduced epithelial transmigration, they retained the capacity to colonize and invade epithelial cells. The *Δfhs* but not the *ΔproABC* mutant was less pro-inflammatory. Hence pneumococcal microinvasion and immune activation are not necessarily pre-cursors to disease progression, highlighting the complex impact of biosynthesis gene mutations on pneumococcal virulence, with differential effects on epithelial colonisation and disease.

## RESULTS

### Deletion of the *proABC* and *fhs* genes have broad-ranging differential effects on the *S. pneumoniae* virulence gene expression and metabolic gene pathways when under stress

We have previously shown that Δfhs and ΔproABC showed considerable derangement in global gene transcription when cultured in human serum compared to growth under optimal nutrient conditions [8]. Metabolic analysis suggested that in sera, Δ*fhs* had an impaired stringent response, and both mutants were under increased oxidative stress and had altered lipid profiles [8]. The Δ*proABC* mutation resulted in the accumulation of glycolytic pathway and peptidoglycan synthesis intermediates. To further understand links between disease attenuation, epithelial colonisation and the innate immune response, we extended the transcriptomic analysis of the mutants’ responses to serum stress [8]. This revealed upregulation of virulence factors such as *ply*, *nanA*, *psaA,* those regulating oxidative stress like *SpxB*, *lctO* and *adhE*, and genes involved in purine and carbohydrate metabolism (Supplemental Information Table 1). Phosphotransferase systems, amino nucleotide sugar, fructose and mannose and metabolic pathways were the most significantly enriched pathways in the *ΔproABC* mutant (Table 1). In contrast, competence, purine, pyruvate, metabolic pathways, amino nucleotide sugar pathways and secondary metabolites were the most significantly enriched pathways in the *Δfhs* mutant under stress (Table 1).

**Table 1.** Differentially expressed pathways from single isogenic mutants in human serum compared to WT. 55 genes were selected within competence pathways, amino acid metabolism, virulence factors, oxidative stress pathways and other known genes important for colonisation, for assessment of changes in gene expression following serum stress. *q-value* measures the proportion of false positives incurred (the false discovery rate) when that particular test is significant.

| Pathway | GeneRatio | P-adjust | qvalue |
| --- | --- | --- | --- |
| *Δfhs* vs. WT |  |  |  |
| Fatty acid metabolism/biosynthesis | 10/57 | 1.08E-09 | 6.49E-10 |
| Biosynthesis of secondary metabolites | 26/57 | 2.09E-04 | 1.26E-04 |
| Purine metabolism | 10/57 | 1.34E-03 | 8.06E-04 |
| Quorum sensing | 11/57 | 3.20E-03 | 1.92E-03 |
| Pantothenate and CoA biosynthesis | 5/57 | 3.59E-03 | 2.16E-03 |
| beta-Lactam resistance | 5/57 | 7.59E-03 | 4.57E-03 |
| Propanoate metabolism | 4/57 | 1.51E-02 | 9.09E-03 |
| 2-Oxocarboxylic acid metabolism | 4/57 | 1.51E-02 | 9.09E-03 |
| Metabolic pathways | 40/57 | 1.94E-02 | 1.16E-02 |
| Amino sugar and nucleotide sugar metabolism | 7/57 | 4.22E-02 | 2.54E-02 |
| Pyruvate metabolism | 5/57 | 4.31E-02 | 2.59E-02 |
| *ΔproABC* vs. WT |  |  |  |
| Fatty acid metabolism/biosynthesis | 10/71 | 1.17E-08 | 8.94E-09 |
| Galactose metabolism | 12/71 | 1.25E-04 | 2.47E-08 |
| Phosphotransferase system (PTS) | 13/71 | 2.54E-03 | 1.93E-03 |
| Amino sugar and nucleotide sugar metabolism | 9/71 | 2.89E-02 | 2.20E-02 |

### *S. pneumoniae* 6B disease-attenuated mutants colonise and microinvade nasal epithelium in EHPC

We investigated whether the *ΔproABC/piaA* or Δ*fhs/piaA* mutations affected epithelial association and microinvasion (Fig 1A) using nasal curette samples from EHPC of healthy volunteers [11]. Visualisation of mucosal cells by confocal microscopy showed colonisation by the WT and mutant pneumococcal strains at day 6 post challenge (Fig 1B): 4/11 volunteers challenged with the WT, 5/11 volunteers with the *ΔproABC/piaA, and* 4/5 volunteers with the Δ*fhs/piaA*. Microinvasion (intracellular bacteria) was seen in 2/11 volunteers with the WT, 5/11 volunteers with the *ΔproABC/piaA,* and 3/5 volunteers with the Δ*fhs/piaA* (Fig 1C). Together, these data show that compared to WT, there was no attenuation of colonisation or microinvasion with either the *ΔproABC/piaA* or Δ*fhs/piaA* strains in EHPC.

**Fig 1.**
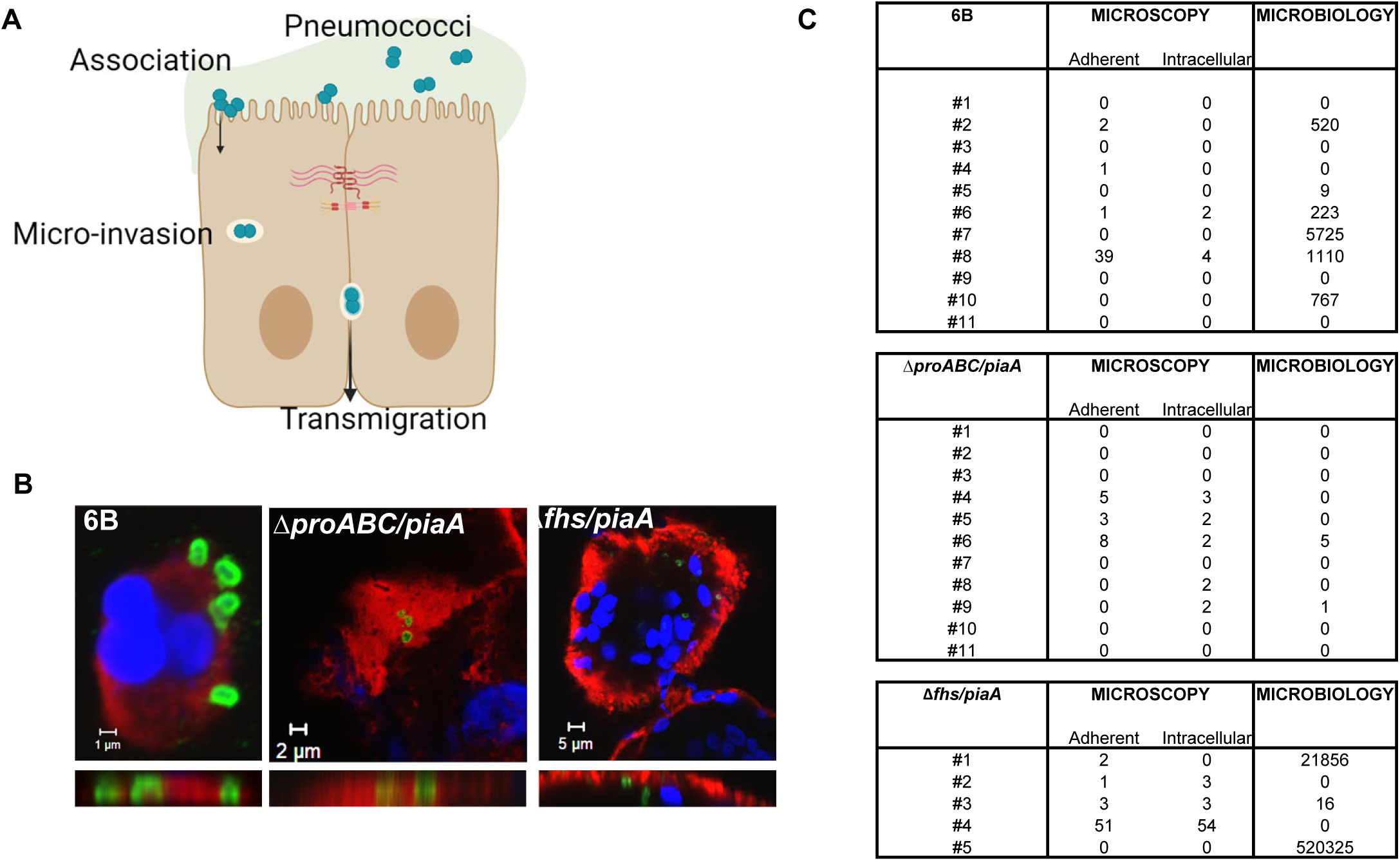
Pneumococcal – epithelial interactions in the EHPC model. (A) Schematic showing that during colonisation, pneumococci adhere to the epithelial cell surface (association), internalise inside epithelial cells (microinvasion). Mucus – lime; pneumococci – purple; intracellular vacuole – dark green; epithelial cells – yellow; nucleus – brown; cell junctions – pink. (B) Representative microscopy images showing pneumococcal-epithelial interactions by the 6B WT, *ΔproABC/piaA* or Δ*fhs/piaA* mutants. Surface carbohydrates (red), nuclei (DAPI, blue), pneumococci (green). (C) Table summarising the microscopy and microbiology counts for each inoculated strain 6 days post challenge per volunteer. Adherent represents pneumococcal surface association and intracellular represents pneumococci that have microinvaded (inside) epithelial cells.

### Colonisation and microinvasion of human airway epithelium by the disease-attenuated mutants is associated with cellular reorganisation and preservation of barrier function

To explore whether disease-attenuation influences epithelial cellular organisation, we infected normal human bronchial epithelial airway (NHBE-A) cells with these pneumococcal strains. We observed epithelial association and microinvasion with all three strains (Fig 2A, top). TEM demonstrated that the intracellular bacteria for all three strains were likely enclosed within vacuoles (Fig 2A, bottom). WT and mutant strains co-localised with Junctional Adhesion Molecule A (JAM-A, Fig 2B, arrow), with no change in JAM-A expression. Quantification by image analysis demonstrated differential effects on the expression of other junctional proteins with increased intensity of Zonula Occludens 1 (ZO-1) at the junctional membrane after infection with WT (Fig 2C), and increased intensity of β catenin staining infected with either *ΔproABC/piaA or Δfhs/piaA*, compared to cells exposed to these strains but not infected (Fig 2D), suggesting cellular reorganisation following pneumococcal exposure.

**Fig 2.**
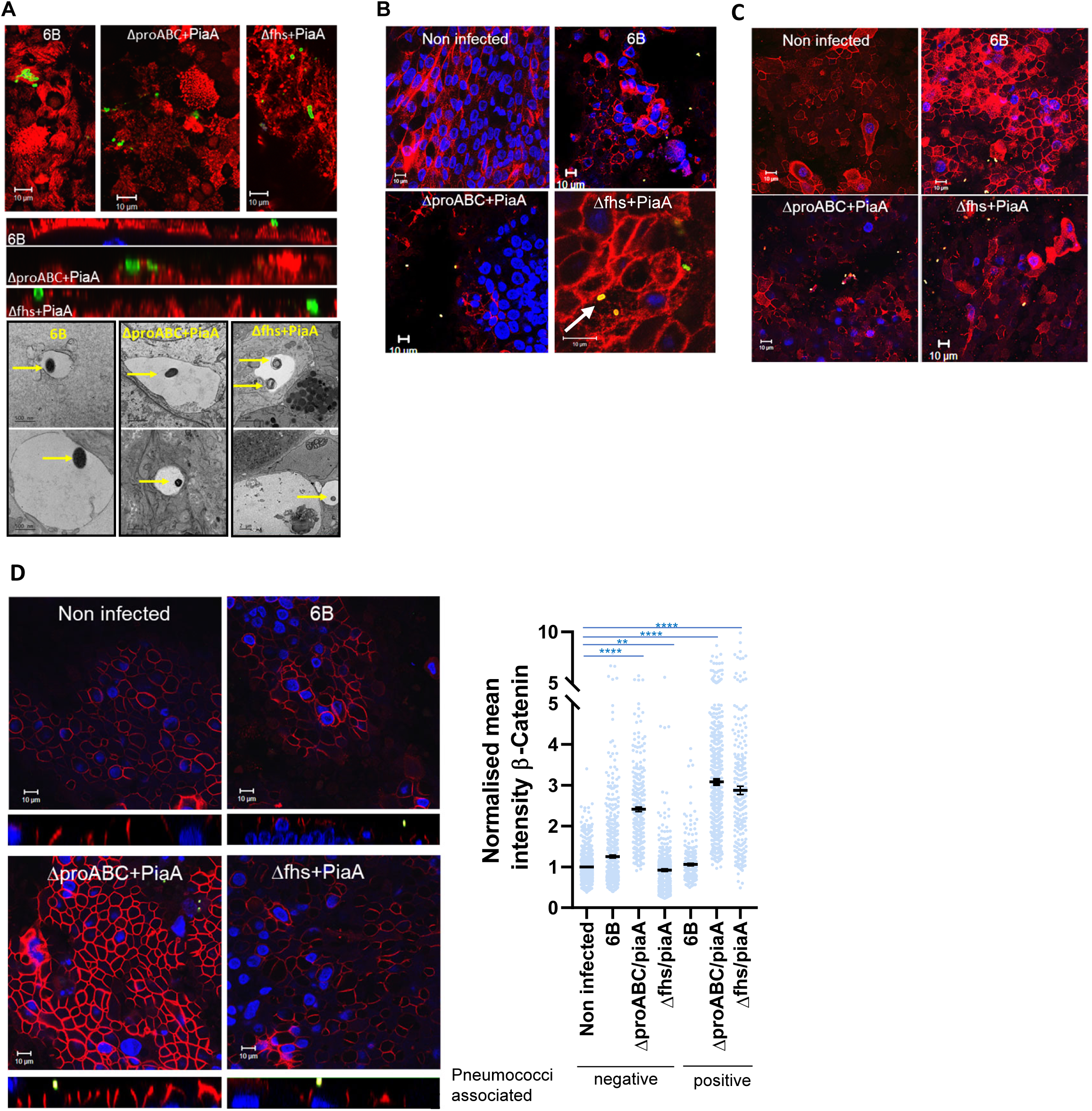
Impact of pneumococcal infection on NHBE-A cell junctional protein expression and barrier integrity. NHBE-A Cells were cultured at ALI and infected with pneumococcal strains for 6hrs. (A) Top: Representative confocal microscopy images from 4 independent experiments with replicates, showing pneumococcal (green) association with the epithelium (red, surface carbohydrates; nuclei blue) at 6hr post infection. Bottom: Representative transmission electron microscopy images from 2 independent experiments, showing pneumococcal microinvasion examples of each strain (arrows), enclosed within a vacuole after 4hr infection. Representative confocal microscopy images from four independent experiments with replicates showing pneumococcal (green) interactions with junctional proteins (red); JAM-A (B), ZO-1 (C) and β catenin (D) in primary HBE-A cells after a 6hr exposure. Nuclei stained with DAPI in blue. Co-localisation of junctional proteins and pneumococci appear yellow, as highlighted by arrows. Integrated intensity of β catenin in pneumococcal negative (not pneumococcal associated) or positive (pneumococcal associated) cells. Non infected v pneumococcal negative cells **** *p* <0.0001 (NI v Δ*proABC/piaA*), ** *p* = 0.0026 (NI v Δ*fhs/piaA*); non-infected v pneumococcal positive cells; **** *p* <0.0001 (NI v Δ*proABC/piaA* and NI v Δ*fhs/piaA*); Kruskal-Wallis, n = 4 independent experiments.

NHBE cells cultured at an air-liquid interface differentiate into a pseudostratified columnar ciliated epithelium, consisting of goblet, Clara, basal and ciliated cells (Fig 3A). Following infection, strains were observed amongst cilia (acetylated tubulin positive cells, Fig 3B, Supplemental Information Fig 1) and the*ΔproABC/piaA* co-localised with mucus (mucin5ac positive, Fig 3C, arrows). Infection with WT and *ΔproABC/piaA* but not *Δfhs/piaA* resulted in an increased abundance of secreted mucus globules compared to uninfected cells (Fig 3A). This appeared due to secretion of preformed mucus rather than de-novo production, as expression of Mucin5AC was unaffected (Fig 3C). Infection with the *ΔproABC/piaA* strain increased expression of acetylated tubulin within pneumococcal-associated cells (Fig 3B), known to be associated with increased microtubule conformation and elasticity [12, 13]. In contrast, only infection with *Δfhs/piaA* caused increased expression of uteroglobin, a multifunctional epithelial secreted protein with anti-inflammatory and anti-chemotactic properties, and a marker for non-ciliated Clara cells (Fig 3D).

**Fig 3.**
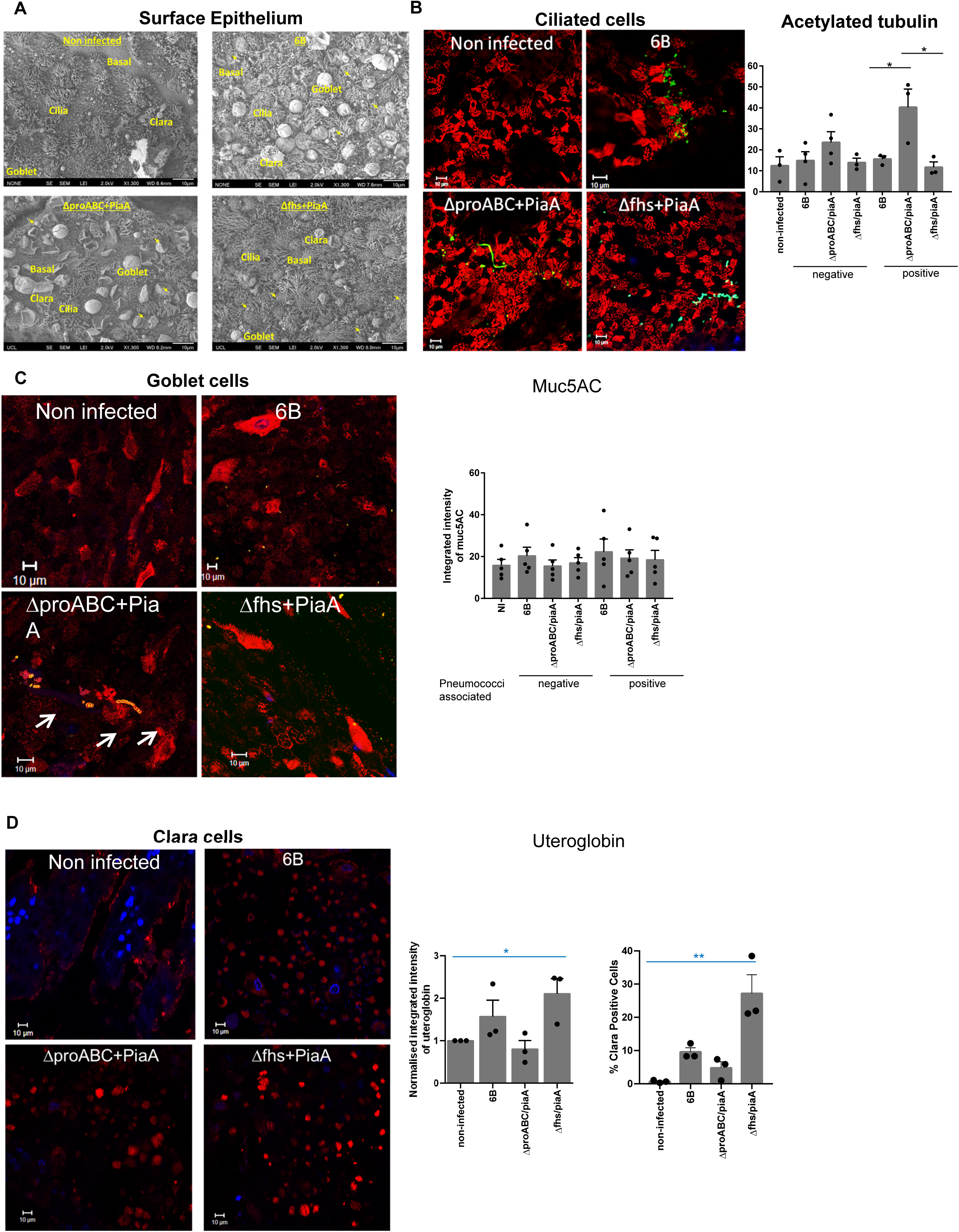
Pneumococcal – epithelial interactions with NHBE-A. Cells were cultured at ALI and infected with pneumococcal strains for 6hrs. (A) Representative scanning electron microscopy images from 2 independent experiments, showing the surface of the epithelium. Representative examples of differentiated epithelia (basal cells, Clara cells, goblet cells, ciliated cells) are shown in yellow text. Colonising pneumococci are highlighted by arrows. Representative confocal microscopy images from 4 independent experiments with replicates for (B) Ciliated cells (acetylated tubulin, red), (C) Goblet cells (mucin5ac, red, (arrows indicate co-localisation of mucin5ac with pneumococci)), (D) Clara cells (uteroglobin, red) after 6hr infection. Pneumococci (green), nuclei (DAPI, blue), colocalization of red and green appears as yellow. Integrated intensity of acetylated tubulin (B) or Muc5ac (C) in pneumococcal negative or positive cells from two or more images. Acetylated tubulin, neg *p* = 0.2677, pos *p* = 0.017, ANOVA. Mucin5ac, neg *p* = 0.6852, pos *p* = 0.8510, N = 3 or 4 independent experiments. (D) Integrated intensity of uteroglobin normalised to non-infected cells (left, *p* = 0.0115 non-infected v infected, Kruskal-Wallis) and percentage of Clara positive cells following infection (right, *p* = 0.0003 non-infected v strains, *p* = 0.0036 6B v mutant strains, Kruskal-Wallis). N = 3 independent experiments.

### *Δfhs/piaA* but not the Δ*proABC/piaA* mutations attenuate the pro-inflammatory epithelial innate response to pneumococcal colonisation and microinvasion

We have shown that Detroit 562 epithelial nasopharyngeal cells are a good model for responses to *S. pneumoniae* [3]. Indeed, infection of Detroit 562 cells showed that WT, *ΔproABC/piaA* and *Δfhs/piaA* strains formed colonies on the epithelial surface (Fig 4A), with limited differences between these strains (Fig 4B, top left). Microinvasion of Detroit 562 cells was observed with all strains (Fig 4B, top right) and showed no differences in intracellular viability (Fig 4B, bottom left) or replication (Supplemental Information Fig 2A-F). In line with attenuation of disease, the *ΔproABC/piaA* and Δ*fhs/piaA* were impaired in their ability to transmigrate across the epithelium (Fig 4B, bottom right). Importantly, the single Δp*iaA* mutation in the *S. pneumoniae* serotype 6B did not affect epithelial association, microinvasion and transmigration, linking the differences seen to the *Δfhs* and *ΔproABC* mutations (Fig 4B). Epithelial β catenin expression was also affected following incubation with the *ΔproABC/piaA* and *Δfhs/piaA* strains, compared to the WT in pneumococcal-associated and non-associated cells. (Fig 4C).

**Fig 4.**
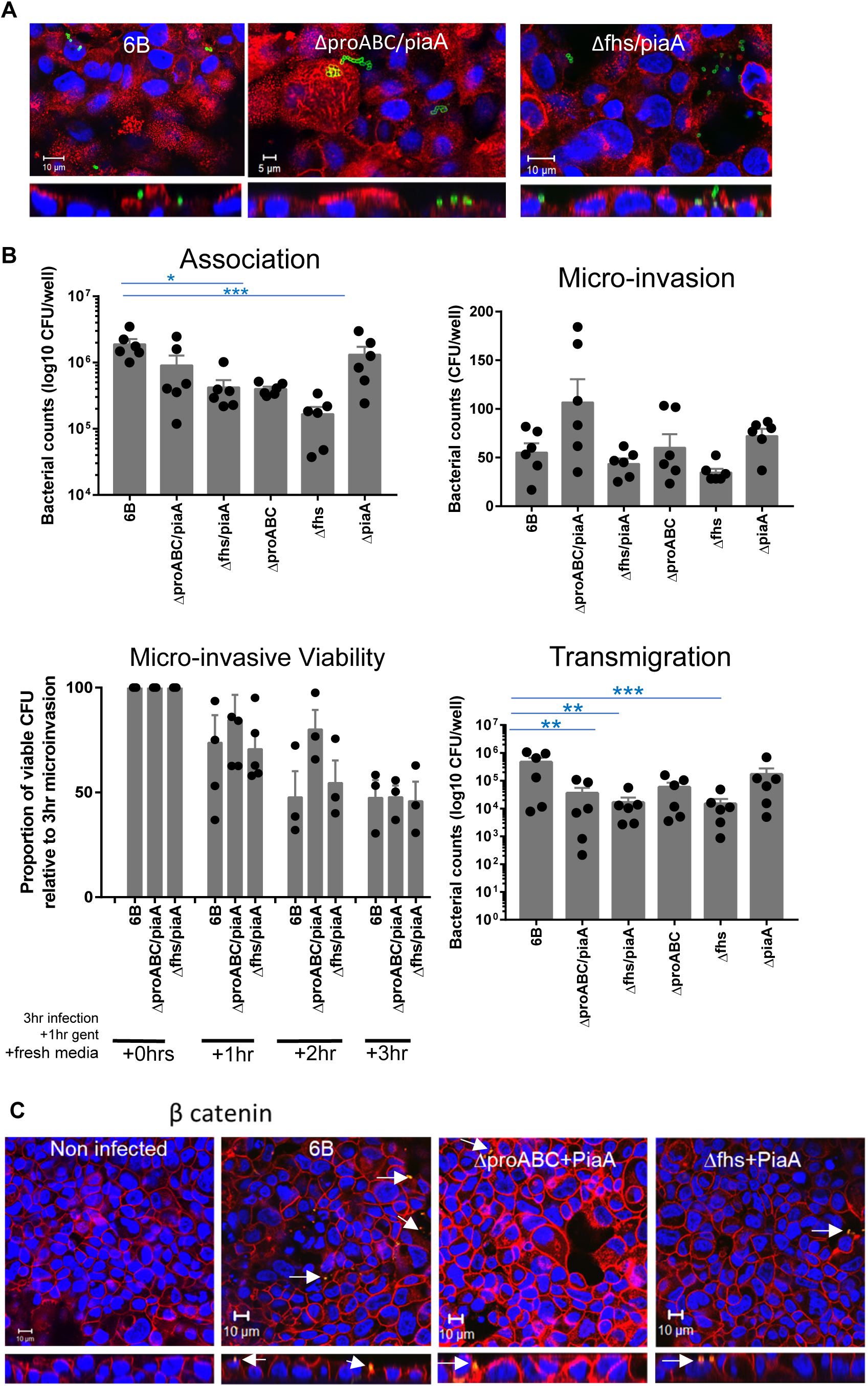
Strain association, microinvasion and transmigration of Detroit 562 cells. Cells were infected with *S. pneumoniae* for three hours and CFU counts recorded. (A) Representative confocal microscopy images from 6 independent experiments with replicates, showing pneumococcal (green) interactions with surface carbohydrates (red). (B, top left) Association of pneumococcal strains to epithelial cells. *p* = 0.0009 (Kruskal-Wallis), n = 6 with replicates. (B, top right) Internalisation of strains into cells, following a gentamicin protection assay. *p* = 0.0219 (Kruskal-Wallis) N = 6 with replicates. (B, bottom left) Pneumococcal viability inside epithelial cells following a gentamicin protection assay. n = 3 or 4 independent experiments with replicates. Counts were normalised to microinvasion counts for each strain after 3 hours infection and compared to 6B WT; +1hr *p* = 0.6517, +2hr *p* = 0.1377, +3hr *p* = 0.1183 (ANOVA). (B, bottom right) Transmigration of pneumococcal strains across confluent monolayers on transwell inserts. CFUs were recorded from the basal chamber. N = 6 independent experiments with replicates. *p* < 0.001 (Kruskal-Wallis). (C) Representative confocal microscopy images from 6 independent experiments with replicates, showing pneumococcal (green) interactions with β catenin (red). Nuclear DAPI stain (blue).

We further assessed links between colonisation, microinvasion and epithelial cell innate immunity using RNAseq. Compared to non-infected cells (Fig 5A), all strains upregulated 78 core epithelial genes related to cytokine signalling in immune system, NLR signalling and GPCR ligand binding. 193 genes were upregulated in response to WT and *ΔproABC/piaA*, characterised by Rhodopsin-like receptors, GPCR ligand binding, cytokine signalling in Immune system, signalling by interleukins and Toll Receptor cascades. Infection with *Δfhs/piaA* resulted in a markedly different transcriptomic profile with modest upregulation of 254 genes, characterised by antigen processing, proteasomal pathways, GTP hydrolysis, and intracellular stress responses (Fig 5A). Overall, the epithelial genes that were most upregulated following infection with WT or *ΔproABC/piaA* were similar and contrasted to the reduced gene expression profile in response to *Δfhs/piaA* despite high levels of cell invasion (Fig 5B). Transcription factors, cellular kinases and cytokines (Fig 5C, 5D) predicted to regulate the differentially expressed genes showed the greatest difference in profile following infection with *Δfhs/piaA* compared to infection with 6B WT and *ΔproABC/piaA*. *Δfhs/piaA* was associated with diminished NFĸB and MAPK activation and instead induced a more intrinsic and epithelial repair response that included TBK1, Raf1, JAK1 kinases and IL-1 and IL-33 cytokine signalling. These data therefore suggest that attenuation of disease-causing potential is not dependent on a reduction in the epithelial innate immune response to colonisation.

**Fig 5.**
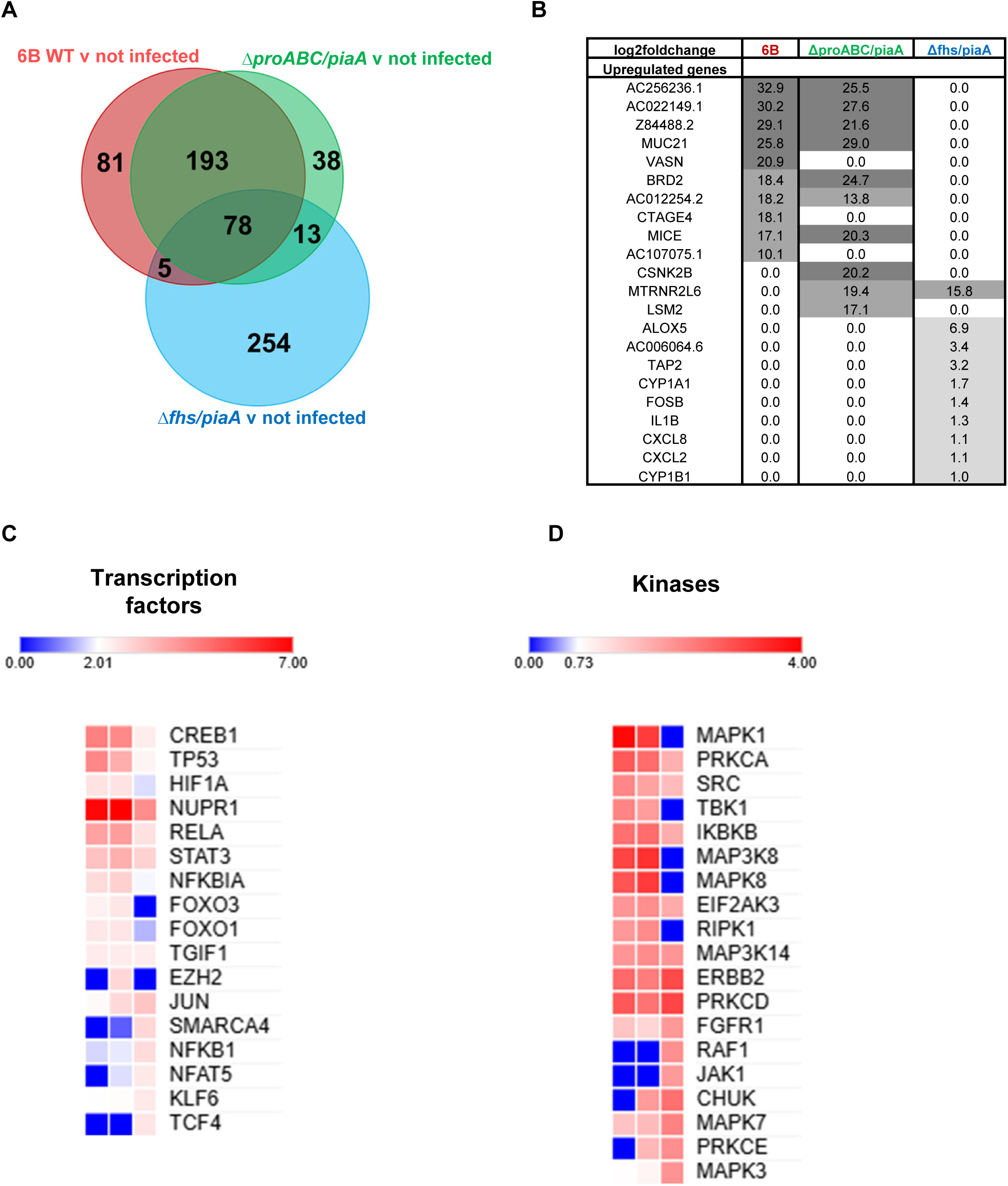
Detroit 562 cell transcriptomic responses to pneumococcal strains. Epithelial cells were infected for four hours with pneumococcal strains and RNA extracted and processed for RNAseq. 6B n = 11, PP n = 17, FP n = 14, non-infected n = 12, from 6 independent experiments. (A) All significantly log2 fold change upregulated genes according to strain compared to non-infected cells. Significance was defined as DEGs with a log2 fold change (logFC) of > 1.5 and false discovery rate (FDR) of < 0.05. (B) Comparison of the top 20 most upregulated genes for each strain, compared to non-infected cells. Upstream regulator analysis for (C) transcription factors and (D) kinases.

### Enhanced epithelial caspase 8 activity following infection with wild-type and Δ*proABC/piaA* strains

To further explore the link between disease attenuation and the epithelial innate immune response to colonisation, we assessed differential caspase activation. The NLRP3 inflammasome drives programmed epithelial cell death via caspase 1, caspase 3/7 and caspase 8 activity [14].

Compared to non-infected cells, all strains increased epithelial caspase activity 6-hour post infection, although this was only significant following infection with *ΔproABC/piaA* (Fig 6A, dark grey bars). Incubating infected cell cultures with either a caspase 1 inhibitor (YVAD-CHO) or caspase 3/7 inhibitor (Z-VAD-FMK), showed that increased caspase activation following infection with WT and *Δfhs/piaA* was not due to caspase 1 activity (Fig 6B)Compared to non-infected cells, caspase 3/7 accounted for 3% and 15% of the caspase activity induced following infection with WT and *Δfhs/piaA* respectively (Fig 6A). In contrast, *ΔproABC/piaA* strain did not induce caspase 3/7 activity. Caspase 8 activation was significantly increased following epithelial infection with 6B WT and *ΔproABC/piaA* strain (Fig 6C).

**Fig 6.**
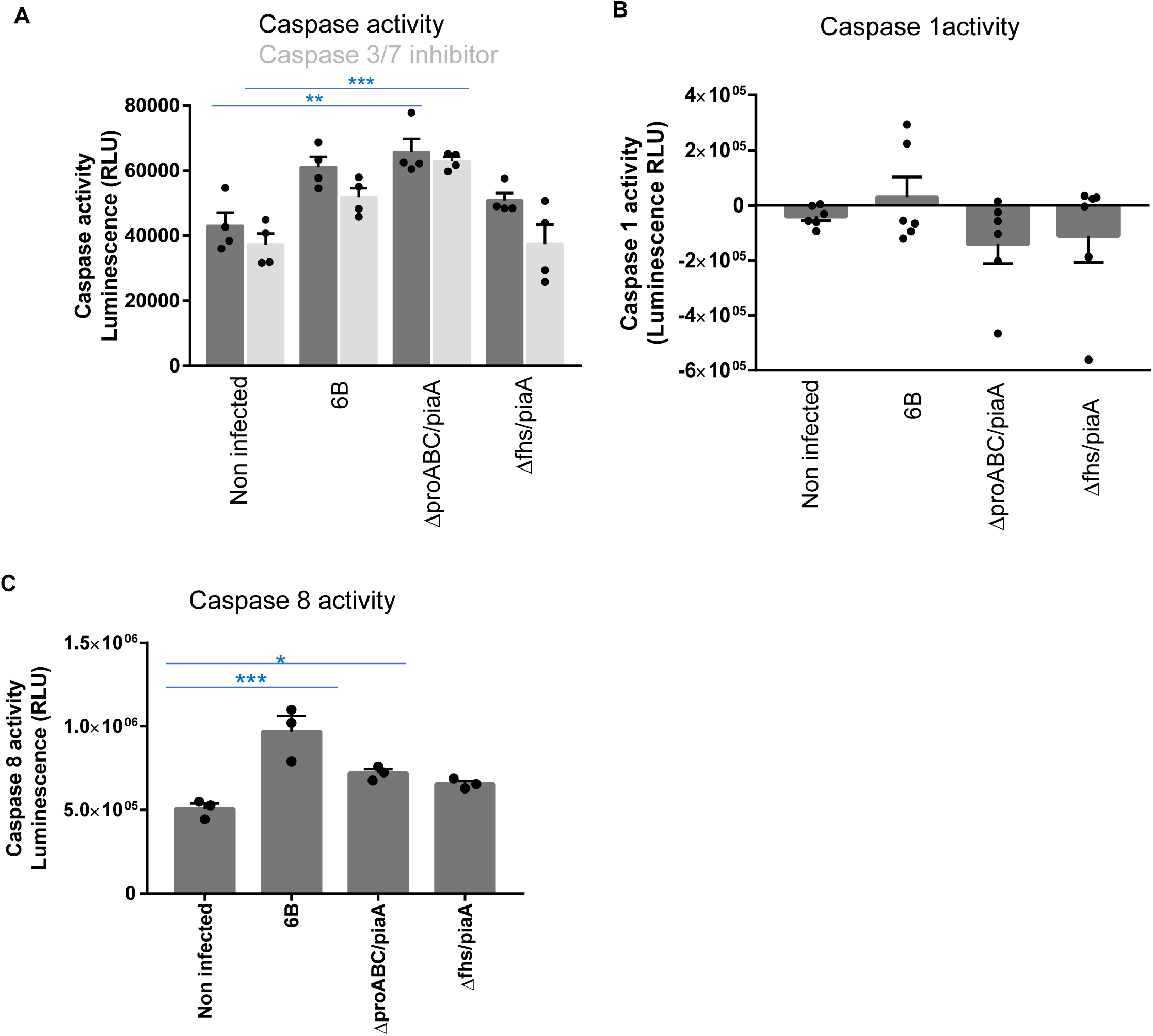
Effects of pneumococcal infection on caspase activity in Detroit 562 cells. Caspase activity was assessed by luminescence, following infection with pneumococcal strains for 6 hours. (A) Caspase 3/7 activity was determined following incubation with the specific inhibitor Z-VAD-FMK. N = 4 with replicates. In comparison to non-infected cells, caspase 3/7 *p* = 0.0022; caspase 3/7 + Z-VAD-FMK inhibitor *p* = 0.0005. In comparison to 6B WT, caspase 3/7 *p* = 0.0215; caspase 3/7 + Z-VAD-FMK inhibitor *p* = 0.0012 (Kruskal-Wallis). (B) Caspase 1 activity was determined following incubation with the specific inhibitor YVAD-CHO. Negative signals indicate caspase activity other than caspase 1. N = 6 with replicates. In comparison to non-infected cells, *p* = 0.39; caspase+YVAD-CHO inhibitor *p* = 0.3604 and, in comparison to 6B WT, *p* = 0.6226; caspase+YVAD-CHO inhibitor *p* = 0.5386 (Kruskal-Wallis). (C) Caspase 8 activity was assessed following infection with pneumococcal strains. N = 3 with replicates. In comparison to non-infected cells, *p* = 0.0015 (ANOVA) and, in comparison to 6B WT, *p* = 0.0172 (ANOVA).

### *fhs* gene mutation is associated with increased pneumococcal hydrogen peroxide secretion and decreased pneumolysin activity

*S. pneumoniae* SpxB and LctO generate hydrogen peroxide (H_2_O_2_) which suppresses ENaC-α transcription and inhibits the Na+–K+ pump, an intracellular signal to increase mitochondrial ROS production [15, 16]. We postulated that differences in cellular reorganisation, the innate inflammatory response and caspase 8 activation seen between *ΔproABC/piaA* and *Δfhs/piaA* mutants could be explained by differential expression of pneumococcal H_2_O_2_. After three and six hours, we detected significantly higher levels of H_2_O_2_ in *Δfhs/piaA* cultures compared to 6B WT and *ΔproABC/piaA* strains (Fig 7A). Furthermore, this was maintained during epithelial cell contact (Fig. 7B).

**Fig 7.**
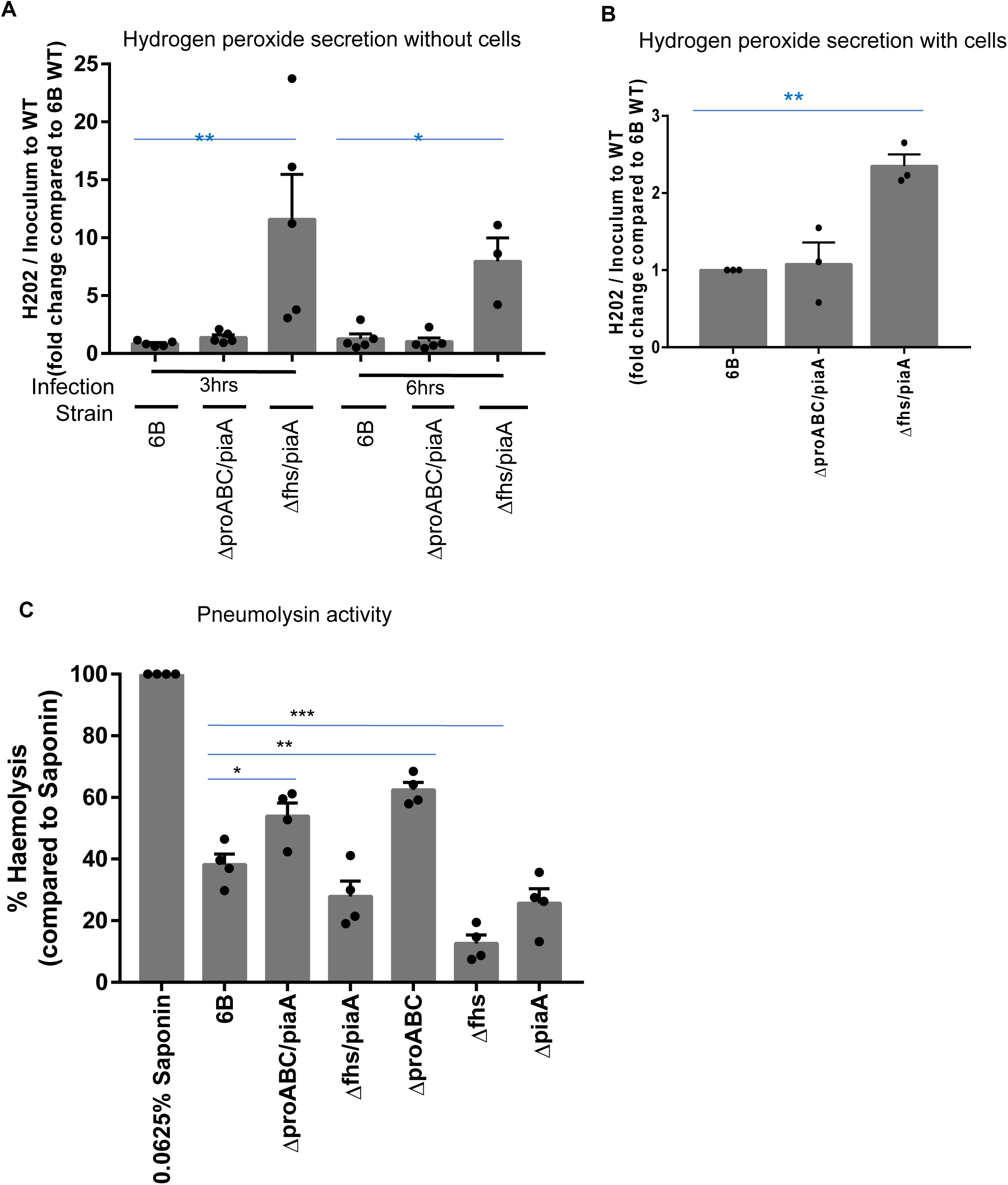
Hydrogen peroxide secretion and pneumolysin activity from pneumococcal strains in the absence and presence of Detroit 562 cells. (A) Fold change of hydrogen peroxide production from the mutants compared to WT, normalised to the CFU counts. 3hr *p* = 0.0087 (ANOVA), 6hr *p* = 0.0209 (Kruskal-Wallis). (B) Fold change of hydrogen peroxide production from the mutants compared to WT, normalised to the CFU counts, following infection of Detroit 562 cells. Media was replaced after 1hr infection and cells incubated for a further 5hrs. N = 3 independent experiments with replicates. Compared to 6B WT, *p* = 0.0034 (ANOVA). (C) Pneumolysin activity was assessed using a red blood cell lysis assay. Strains were incubated with red blood cells for 30 minutes and absorbance measured. N = 4 experiments with replicates. Pneumococcal strain haemolytic activity is calculated in comparison to 100% lysis by 0.0625% saponin. *p* <0.0001 compared to 6B WT (ANOVA).

Pneumolysin can both activate and suppress the host innate immune response and mediates the haemolytic activity characteristic of the pneumococcus [17–19]. H_2_O_2_ increases the release of pneumolysin but paradoxically also negatively impairs its haemolytic activity [20]. In line with this paradoxical effect the higher production of H_2_O_2_ by the *Δfhs/piaA* strain, was associated with increased *ply* gene expression but reduced red blood cell lysis compared to 6B WT and *ΔproABC/PiaA* (Fig. 7C).

## DISCUSSION

We show that when using isogenic serotype 6B biosynthesis pathway mutant strains, attenuated for systemic disease but not for carriage, epithelial microinvasion is preserved [5, 8, 21]. Epithelial barrier transmigration was significantly less with the isogenic strains with the Δ*fhs/piaA* and Δ*proABC/piaA* mutations compared to WT *S. pneumoniae* 6B. Importantly, while epithelial microinvasion, cellular reorganization, the innate epithelial immune response, and the propensity to cause invasive disease following pneumococcal colonisation are all strain-dependent, no single determinant explained these differences.

In our primary epithelial cell infection model [22], consisting of Clara cells (involved in secretory, xenobiotic metabolic, and progenitor cell functions), goblet cells (mucus production), ciliated cells (small particle transport), infection with WT and mutant strains resulted in epithelial cellular re-organisation involving β catenin, acetylated tubulin and uteroglobin. However, strain differences seen on the impact on junctional or cytoskeletal protein organisation or mucus production did not explain the disease attenuation seen with Δ*fhs/piaA* and Δ*proABC/piaA*.

Epithelial infection with WT and *ΔproABC/piaA* induced innate inflammatory pathway gene expression [3], including cytokine/chemokines, interleukins, Toll Like Receptor and NFĸB signalling, alongside apoptotic transcripts such as FOXO3 and TP53. Notably, some epithelial genes were upregulated exclusively during WT infection, which could be related to its greater disease-causing potential. These were involved in RNA degradation, O-linked glycosylation, wnt signalling, DNA sensing, RIG-I pathways all of which play a role in the immune response and signal transduction. In contrast, the *Δfhs/piaA* strain activated a distinct pro-survival and adaptation profile, upregulating genes such as JUN, SMARCA4, NFAT5, KLF6 and TCF4 which are involved in homeostatic signalling and chromatin remodelling [23, 24], and suggesting a more tolerant epithelial response to this strain. Ultimately, these capsule-independent differences in pneumococcal-epithelial interactions may shape the differences seen in adaptive immune response identified using EHPC to colonisation with the WT, *ΔproABC/piaA* or *Δfhs/piaA* strains [11, 25].

Importantly, while all strains adhered and microinvade the epithelium, WT serotype 6B infection led to pneumococcal transmigration, a more pro-inflammatory innate immune transcriptomic profile and increased caspase 8 activity, indicative of a robust cellular response to *S. pneumoniae*, with Δ*proABC/piaA* and *Δfhs/piaA* exhibiting attenuation of these responses. This suggests that pneumococcal sensing by the epithelium is affected by disruption of *S. pneumoniae* metabolism [18, 26]. Pneumolysin is a key driver of epithelial inflammatory responses and increased acetylated tubulin and microtubule stabilisation [13] [3]. Under stress conditions, the Δ*proABC* and Δ*fhs* mutations upregulated multiple metabolic pathways, presumably as a compensatory mechanism, and also genes encoding several pneumococcal virulence factor genes (Graphical Abstract Summary) including pneumolysin and H_2_O_2_. Based on these findings, we propose that differential expression of pneumolysin, and H_2_O_2_ by the WT and mutant strains may contribute to the distinct patterns of epithelial responses that we observed, with the potential to influence both bacterial clearance and transmission [1]. Whether the distinct epithelial innate immune response to *Δfhs/piaA* infection is due to diminished NFĸB and MAPK activation (potentially linked to extracellular receptor engagement), reduced caspase 8 activity mediated by enhanced pneumococcal H_2_O_2_ release, decreased pneumolysin activity, or a compensatory metabolic response to the stress, remains to be determined. As seen in ‘professional’ intracellular bacterial species such as *Listeria* and *Salmonella*, the host response to microinvasion might also promote pneumococcal transmission [27, 28]. Whether the less pro-inflammatory *Δfhs/piaA* is less transmissible than the more inflammatory WT is unknown, which could be explored using the EHPC model [29].

In conclusion, these data further support our proposed paradigm that epithelial microinvasion by *S. pneumoniae* is frequent and does not necessarily lead to disease. Microinvasion may aid transmission through immune evasion and localised inflammation and, transmigration across the epithelium may represent the first step towards transition to invasive disease. Our findings also highlight the broad-ranging effects of single gene mutations in pneumococcal biosynthesis pathways, which may explain the complexity of differences between seemingly closely related strains. The interplay between *S. pneumoniae* and the host epithelium is modulated by key mediators such as hydrogen peroxide and pneumolysin, which can have both enhancing and inhibitory effects on epithelial immune responses depending on the infection context. However, strain differences in subsequent epithelial cell microinvasion, cellular reorganisation and inflammatory responses appear driven by the differential expression of multiple bacterial virulence and metabolic pathways, rather than any single determinant.

## METHODS AND PROTOCOLS

### Bacteria

The *Streptococcus pneumoniae* serotype 6B (strain BHN 418 [30]) was a kind gift from Prof. Birgitta Henriques Normark (Karolinska Institute). *ΔproABC/piaA* and *Δfhs/piaA* mutant strains were generated using overlap extension PCR as described by Ramos-Sevillano *et al* [5]. Aliquot stocks frozen at 0.3 O.D._600nm_ were defrosted, centrifuged at 8000g for eight minutes and resuspended in 1% Foetal Bovine Serum (FBS, Gibco) in MEM media (Gibco) for infections. Starting inoculums were not significantly different across strains for all assays.

### Bacterial RNA-seq sample collection, sequencing and reads

Pneumococcal strains were grown in THY to mid-log phase and pellets resuspended in THY and undiluted human serum for 60mins. The culture was centrifuged and stored in RNA Protect at −70°C before extracting total RNA [8]. Total RNA from each library was ribo-depleted and sent for Illumina Next Seq Sequencing Pathogen Genomics Unit, UCL, as detailed in Supplemental Information.

### Differential gene expression analyses

The generated count matrix was imported into R-studio (R v3.4.2) and normalization of counts and differential gene expression analysis was performed by DESeq2 [31]. DESeq2 normalized libraries were regularized log transformed for visualizing heatmaps and clustering. Differential gene expression was performed on raw counts. Fold change shrinkage was implemented with the apeglm package within DESeq2. Genes with a log2 fold change > 1.5 and false discovery rate (FDR) of < 0.05 were categorized as differentially expressed. KEGG pathway enrichment and module analysis were performed on differentially regulated genes using over representation analysis (ORA) implemented by clusterProfiler [32].

### Experimental Human Pneumococcal Carriage Model (EHPC)

A full explanation of experimental design and selection criteria has been previously described [3] and detailed in Supplemental Information.

### Normal Human Epithelial Cells

Normal Human Bronchial/Tracheal Epithelial Cells (NHBE-A, ATCC® PCS-300-010™) were plated at 6×10^3^ cells/cm^2^ on a layer of sub-lethally γ-irradiated (60 Gy) mouse embryonic fibroblasts 3T3-J2 cells (kind donation from Howard Green, Harvard Medical School, Boston [33]) with keratinocyte culture medium cFAD (3:1 DMEM (Gibco) to Ham F-12 Nut Mix (Gibco), 10% FBS (HyClone, Sigma), 1% Penicillin-Streptomycin (100X, Sigma), 0.4 μg/mL Hydrocortisone (Calbiochem, USA), 5 μg/ml Insulin (various), 10-10 Cholera Toxin (Sigma) and 2×10-9 Triodothyronine (Sigma)). Epithelial cells were stimulated with 10 ng/mL human epidermal growth factor (hEGF, PeproTech, USA) on days 3 and 5. Cultures were grown at 37°C 6% CO_2_. For NHBE-A differentiation in Air Liquid Interface culture, cells were plated at 6×10^4^/cm^2^ on 1μm pore PET cell culture inserts (Greiner). All chambers were supplemented with cFAD medium. After 48 hours, medium was removed from the top chamber. Lower chamber media was replaced with PneumaCult™ differentiation Medium (Stemcell Technologies) every 2 days for 3 weeks. Antibiotics were removed from media and cells were washed 24 hours before infection experiments.

### Human Epithelial Cell Lines

Human pharyngeal carcinoma Detroit 562 epithelial cells (ATCC_CCL-138) were grown in 10% FCS in alpha MEM media (Gibco). For each set of experiments, cells were used within 5 passages. Cells consistently tested PCR negative for mycoplasma (Cambridge Biosciences).

### Pneumococcal-epithelial cell co-culture

For association and internalisation assays, confluent monolayers of Detroit 562 cells cultured on 12 well plates (Corning), were infected(1 cell to 10 pneumococci) for three hours in 1% FCS MEM or six hours for HNBE-A cells in 1% Pneumocult media without supplements [3]. Cells washed three times in HBSS^+/+^ before incubation in 1% saponin for 10 minutes at 37°C. Bacterial dilutions were plated on horse blood agar for 16 hours and colony forming units counted. For internalisation quantification, 200µg/ml gentamicin was added one hour before incubation with saponin.

For transmigration assays, confluent monolayers of Detroit 562 cells cultured on 3µm pore, PET Transwell Inserts (ThermoFisher) [3] were infected for three hours. 50µl of basal media was repeatedly removed to measure pneumococcal load by counting CFU/well.

### Confocal Microscopy

Epithelial monolayers on transwell membranes were infected then fixed in 4% PFA (Pierce, Methanol Free) as previously described and detailed in Supplemental Information [3]. Cells were processed for image analysis using an inverted Zeiss LSM (either 700 or 880). Z stacks were recorded at 1µm intervals at either 40x oil or 63x oil objectives. Automated image analysis was carried out using Cellprofiler [34].

### Electron Microscopy

Preparation of Detroit 562 cells and NHBE-A cells was performed as previously described and details can be found in Supplemental Information [3].

### Pneumolysin activity

Adapted from Kirkham *et al* [35]. Final concentration of 2% horse blood (EO Labs) solution was prepared in phenol free alpha MEM (Life Technologies) and added to a 96 U-bottom plate (Corning). 2.5×10^6^ bacteria were added to 96 well. Saponin was used as a positive control for lysis. Solutions were incubated for 30 minutes at 37°C before centrifugation at 1000g for one minute. Solution was transferred into a new 96 well plate and absorbance recorded at 540nm on a plate reader.

### Hydrogen Peroxide Assay

Detroit 562 cells were cultured in white 96 well plates (Corning). Cells were incubated with H_2_O_2_ substrate and infected for up to six hours according to manufacturers’ instructions (ROS-Glo^TM^ H_2_O_2_ Assay, Promega). Luminescence was read using a GloMax-Multi Detection System plate reader (Promega).

### Caspase assay

Detroit 562 cells were cultured in white 96 well plates (Corning). Cells were infected for up to six hours. Primary cell supernatant was transferred to a white 96 well plate. Samples were incubated with Caspase-Glo® 1 Reagent, Caspase-Glo® 1 YVAD-CHO Reagent, Caspase-Glo® Z-VAD-FMK Reagent or Caspase-Glo® 8 Reagent, for one hour and according to manufacturers’ instructions (Promega). Luminescence was read using a GloMax-Multi Detection System plate reader (Promega).

### RNA samples and sequencing (RNA-Seq)

Confluent monolayers of Detroit 562 epithelial cells were infected for four hours before collection in RNALater. RNA was extracted using a Qiagen RNAEasy micro kit and treated for DNA removal using a Qiagen Turbo DNA-free kit. RNA quality was assessed and quantified using a BioAnalyser (Agilent 2100). Samples were sequenced using Illumina Nextseq 500/550 High Output 75 cycle kit giving 15-20 million 41bp paired-end reads per sample (Pathogens Genomic Unit, UCL).

### Statistics

Conditions within each experiment were performed in duplicate or triplicate, and each experiment was performed independently more than three times, unless stated otherwise. Error bars represent SEM, unless stated otherwise. GraphPad Prism Version 7 was used for parametric (t-tests or ANOVA) or non-parametric (Mann-Whitney or Kruskal-Wallis tests) analysis, which was based on the Shapiro-Wilk normality test. Ad hoc tests were performed using Tukey’s (parametric data) or Dunn’s (non-parametric data) multiple comparisons test. P values <0.05 were considered significant.

### Data and code availability

Source data are provided as a Source Data file. RNAseq data from the Detroit 562 cell infections is publicly accessible through Zenodo with Accession (DOI:10.5281/zenodo.7997789).

## ACKNOWLEDGEMENTS

This study was funded by CMW from a Postdoctoral Innovation Award from UCL, a Pump-Priming Award and Enhancement Award from the Human Infection Challenge Network for Vaccine Development (HIC-VAC (funded by the GCRF Networks in Vaccines Research and Development) which is co-funded by the MRC and BBSRC). This UK funded award is part of the EDCTP2 programme supported by the European Union. CMW and RSH are supported by the Medical Research Council (grant MR/T016329/1). The authors would like to thank Prof. Peter Hermans (Radboud University Medical Centre) for donating the 6B pneumococcal strain originally from Prof. Birgitta Henriques-Normark (Karolinska Institute) and obtained directly from DF at LSTM. EHPC sample collection was supported by the Medical Research Council (grant MR/M011569/1), Bill and Melinda Gates Foundation (grant OPP1117728) awarded to DF and the National Institute for Health Research (NIHR) Local Comprehensive Research Network. The authors also wish to thank the EHPC Clinical Team at LSTM and all the volunteers. Confocal imaging facilities at LSTM were funded by a Wellcome Trust Multi-User Equipment Grant (104936/Z/14/Z). Electron microscopy processing and image capture was performed by Dr. Elizabeth Slavik-Smith in the Biosciences EM facility at UCL. MC is supported by a Springboard to Independence grant (AirwayStasis) from the French Government’s Investissement d’Avenir program, the Laboratoire d’Excellence ‘‘Integrative Biology of Emerging Infectious Diseases” (ANR-10-LABX-62-IBEID) and the Chromatin and infection unit headed by Melanie A. Hamon. RNA-Seq library preparation was undertaken at UCL through the UCL/UCLH Biomedical Research Centre and MRC funded Pathogen Genomics Unit (PGU, G0900950). All RNAseq data processing was completed by the PGU. MN is supported by the Wellcome Trust and by NIHR Biomedical Research Centre Funding to University College Hospitals NHS Foundation Trust and University College London. RR is supported by Marie Skłodowska-Curie Individual Fellowships 896014. PB is supported by the European Research Council (ERC-Stg No. 639429), the Rosetrees Trust (M362-F1; M553), the CF Trust (SRC006;SRC020), the London Advanced Therapies - Research England (C2N-AT.006), the MRC Confidence in Concept scheme (Award: MC_PC_17180), the Innovate UK (Smart grant n.10005465), the Duchenne Parent Project (DPP) and the NIHR GOSH BRC. MW is supported by a 223065 Wellcome Investigator Award to Clare Jolly, UCL. ERS and JSB are supported by MRC grant R/N02687X/1 and Wellcome grant 221803/Z/20/Z. JSB, MN, and RSH acknowledge funding from the Department of Health’s National Institute of Health and Care Research Biomedical Research Centre funding to UCL and UCLH. RSH is a NIHR Senior Investigator.

## AUTHOR CONTRIBUTIONS

CMW conceived, funded, designed and conducted the study, performed the experiments, acquired samples and processed data, and wrote the first draft of the manuscript. GP analysed and interpreted the RNAseq data. MB analysed and interpreted the bacterial transcriptomic data. RR cultured and differentiated primary epithelial cells. ERS generated the 6B mutant strains and bacterial transcriptomic data. JR collected and processed EHPC samples. JAGA processed the RNA-Seq data. MW ran the image analysis. MC provided technical advice. PB and CJ provided investigator support. MN provided RNA-Seq analysis support. DF provided investigator support and EHPC clinical samples. JSB generated the 6B mutant strains, interpreted the data and revised the manuscript. RSH provided investigator support, designed and interpreted the data, revised the manuscript. All authors commented on and approved the manuscript.

## COMPETING INTERESTS

The authors declare no competing interests.

## Notes

### Competing Interest Statement

The authors have declared no competing interest.

### Summary of Updates

The manuscript has been altered in terms of narrative, order, length and figure generation.

