## Supplemental Information for "Disease-attenuated pneumococcal biosynthesis gene mutants invade the mucosal epithelium and induce innate immunity"

### SUPPLEMENTAL METHODS AND PROTOCOLS

#### Bacterial RNA-seq sample collection, sequencing and reads

Mutant pneumococcal strains (three biological replicates for each mutant) and 6B WT (three biological replicates) were grown in THY to mid log phase and the pellets resuspended in THY and undiluted human serum for 60mins. The culture was then pelleted, and RNA stored in RNA Protect at -70°C before extracting total RNA [1]. Total RNA from each library was ribo-depleted and sent for Illumina Next Seq Sequencing Pathogen Genomics Unit (PGU), UCL. The quality of raw FASTQ reads was checked by FastQC v0.11.5, Babraham Bioinformatics, UK and visualised using multiQC v1.9 [2]. Reads were trimmed using Trimmomatic 0.39 [3]. Trimmed reads were also checked by FastQC and multiQC. Trimmed reads were then mapped to KEGG annotated 6B genome sequence (670-6B, Accession: CP002176.1) using bowtie2 with default settings [4]. Conversion into BAM files was performed using SAMtools [5]. Mapped reads were visualized in the Integrated Genome Browser [6]. FeatureCounts v2.0.0 was used to summarize read counts for each annotated feature in the respective reference genomes in multimapping mode (-M) [7].

#### Experimental Human Pneumococcal Carriage Model (EHPC)

A full explanation of experimental design and selection criteria has been previously described [8]. Following written informed consent, healthy non-smoking adults between the ages of 18 – 59 were inoculated with 80,000 CFU/nostril live strains of *S. pneumoniae*. Mucosal cells from the inferior turbinate using a plastic Rhino-probe™ (curettage, Arlington Scientific) were collected prior to inoculation, and days 2 and 6 post-inoculation. Samples were processed for confocal microscopy analysis. Nasal wash samples were collected for CFU count analysis. There were no adverse events. A positive carrier was defined as colonisation at any time point (day 2 of day 6 post inoculation) as detected by culture [8]. Ethical approval was given by NHS Research and Ethics Committee (REC)/Liverpool School of Tropical Medicine (LSTM) REC, reference number: 18/NW/0481 and Human Tissue Authority licensing number 12548.

**Confocal Microscopy**

Sample processing was performed as previously described [8]. Briefly, mucosal cells from the EHPC model were placed directly into 4% PFA and cytospun onto microscope slides. NHBE-A cells and Detroit 562 cells cultured on transwell membranes were fixed in 4% PFA (Pierce, Methanol Free). Cells were permeabilised with 0.2% Triton X-100 for 10 minutes, blocked for one hour in blocking buffer (3% goat serum and 3% BSA in PBS (Sigma)) and incubated with pneumococcal antisera Pool Q (raised against the polysaccharide capsule, SSI Diagnostica) for one hour. Wheat germ agglutinin conjugated to Rhodamine (Vector Labs) was added along with goat-anti rabbit IgG conjugated to Alexa Fluor 488 for 45 minutes in blocking buffer. Primary (used at 1:100) antibodies used for epithelial cell proteins can be found in Supplemental Information Table 2. Secondary (used at 1:500) antibodies were either goat anti-mouse or goat anti-rabbit Alexa Fluor 555 or 546, ThermoFisher. Of note, for epithelial protein antibodies raised in rabbit (Uteroglobin), no pneumococcal staining was possible and therefore pneumococci were not visible on the images. DAPI was added for 5 minutes and cells were mounted with Aqua PolyMount (VWR International) and a coverslip on the microslide. Pneumococci were recorded by manual scanning of the whole cytospin for the EHPC samples. More than three fields of view were recorded for each bacterial strain for Detroit 562 cell infections, using inverted Zeiss LSM (either 700 or 880) confocal microscopes. Images were processed with Zeiss LSM Image Browser. Z stacks were recorded at 1μm intervals at either 40x oil or 63x oil objectives. Automated image analysis was carried out using Cellprofiler [9].

**Electron Microscopy**

Preparation of Detroit 562 cells and NHBE-A cells was identical and previously described [8]. Briefly, cells cultured on transwells were fixed with 2% paraformaldehyde and 1.5% glutaraldehyde in 0.1 M cacodylate buffer and post-fixed in 1% OsO4 / 1.5% K4Fe(CN)6 in 0.1 M cacodylate buffer pH7.3. For scanning electron microscopy, samples were dehydrated in graded ethanol-water series and infiltrated with Agar 100 resin mix. Ultra-thin sections were cut at 70-80 nm using a diamond knife on a Reichert ultracut microtome. Sections were collected on 300 mesh copper grids and stained with lead citrate. For transmission electron microscopy the samples were hardened at 60$^{o}$C for 48 hours. The coverslip was removed, and a representative area was selected. Ultra-thin sections were cut at 70-80 nm using a diamond knife on a Reichert ultra-cut S microtome. Sections were collected on copper grids and stained with lead citrate. Samples were viewed with a Joel 1010 transition electron microscope and the images recorded using a Gatan Orius CCD camera.

**TABLE AND FIGURE LEGENDS**

**Table 1. Differentially upregulated genes by *S. pneumoniae* biosynthesis mutants in comparison WT under serum stress.** Enrichment analysis of virulence genes, experimentally deleted biosynthesis genes, purine related genes, hydrogen peroxide regulators and carbohydrate metabolism. A comparison of differential gene expression was assessed when pneumococcal strains were cultured in human serum, comparing the single ∆*proABC* ∆*fhs* mutant relative to the WT strain.

**Table 2. Primary antibodies used for Microscopy.** Primary and secondary antibodies for epithelial proteins. Primary antibodies were added for 1 hr at room temperature before washing off in HBSS and adding secondary antibodies for 45 minutes.

**Fig 1. Ciliated Primary HBE-A cells.** Representative example of pneumococcal (green) associations with ciliated (acetylated tubulin, red) cells as illustrated with the Δ*proABC+PiaA* mutant in primary HBE-A cells.

**Fig 2. Association, microinvasion and transmigration of pneumococci following 1hr infection in Detroit 562 cells.** Pneumococcal-epithelial interactions at 1hr post infection. Association, micro-invasion and transmigration of pneumococcal strains after 1hr infection in Detroit 562 cells and CFUs recorded. (A) Association of 6B strains. $p = 0.0024$. (B) Internalisation of 6B strains. $p = 0.0107$. (C) Transmigration of 6B strains after 1hr infection. Pneumococcal strains were added to the apical chamber of a transwell insert with a confluent monolayer of Detroit 562 cells. CFUs were recorded over time from the basal chamber, reflecting bacteria that had transmigrated across the epithelium. $p = 0.0106$. (D) Pre ($p = 0.5403$) and post ($p = 0.6384$) inoculum. N = 4 independent experiments with replicates (Kruskal-Wallis). (E) Pre and post inoculum after 3 hours infection. Pre- (light grey) and post- (dark grey) inoculums after 3 hours incubation with epithelial cells. Pre- $p = 0.8366$, Post- $p = 0.0160$, 6B v isogenic mutants (Kruskal-Wallis), n = 6 with replicates. (F) Cell supernatant from infected cells was analysed for lactate dehydrogenase secretion. N = 3 - 5 with replicates. $p = 0.046$ compared to non-infected cells, and $p = 0.5649$ between 6B WT and mutant strains (ANOVA).

**Supplemental Table 1:** Differentially upregulated genes by *S. pneumoniae* biosynthesis mutants in comparison WT under serum stress

| Gene | Δ*proABC* vs. WT | | Δ*fhs* vs. WT | |
|---|---|---|---|---|
| | FoldChange | P-adjust | FoldChange | P-adjust |
| | | Deleted biosynthesis genes | | |
| *fhs* | 1.16 | 3.30E-03 | -12.04 | 3.10E-54 |
| *proA* | -11.31 | 2.39E-37 | 1.38 | 1.06E-02 |
| *proB* | -11.47 | 8.22E-34 | 1.35 | 1.59E-02 |
| *proC* | -11.47 | 1.74E-27 | 1.38 | 4.84E-02 |
| | | Virulence Genes | | |
| *nanA* | 7.75 | 9.88E-20 | 2.87 | 1.92E-11 |
| *nanB* | -1.03 | 9.24E-01 | -1.36 | 1.39E-01 |
| *ply* | 5.91 | 6.74E-33 | 9.22 | 9.05E-94 |
| *psaA* | 2.93 | 4.21E-15 | 3.73 | 3.81E-41 |
| | | Purine metabolism | | |
| *purN* | 3.33 | 1.89E-08 | 6.40 | 9.12E-26 |
| *SP670_0127* | 1.99 | 8.51E-03 | 2.96 | 8.95E-07 |
| *purH* | 1.84 | 3.77E-03 | 3.81 | 7.48E-14 |
| *purD* | 2.80 | 4.33E-08 | 3.22 | 6.10E-14 |
| *purE* | 2.23 | 4.27E-04 | 2.67 | 8.14E-07 |
| *purK* | 3.27 | 6.15E-09 | 3.61 | 2.62E-16 |
| *purB* | 2.03 | 4.67E-05 | 2.11 | 2.91E-09 |
| *SP670_0123* | 3.66 | 2.48E-18 | 6.96 | 3.83E-48 |
| | | Oxidative stress | | |
| *spxB* | 1.12 | 5.65E-01 | 1.01 | 9.43E-01 |
| *lctO* | 3.17 | 4.16E-13 | 1.39 | 1.02E-02 |
| *spx* | -1.03 | 9.16E-01 | -1.45 | 4.84E-02 |
| *adhE* | 1.32 | 1.83E-01 | 1.31 | 5.90E-02 |
| *dpr* | -1.30 | 0.14 | -1.88 | 5.79E-06 |
| | | PTS/ Carbohydrate metabolism | | |
| *SP670_0137* | 64.83 | 1.93E-60 | 1.69 | 3.82E-04 |
| *SP670_0138* | 67.98 | 4.63E-51 | 2.00 | 6.16E-04 |
| *SP670_0139* | 43.35 | 6.09E-44 | 1.67 | 1.88E-03 |
| *SP670_0140* | 41.35 | 1.64E-14 | 1.84 | 1.75E-03 |
| *SP670_0141* | 33.44 | 1.73E-12 | 1.93 | 3.36E-04 |
| *SP670_0142* | 22.86 | 2.26E-10 | 1.77 | 8.37E-05 |
| *SP670_0700* | 36.80 | 1.32E-11 | 1.98 | 2.07E-02 |
| *SP670_0701* | 42.56 | 5.29E-55 | 1.67 | 1.40E-02 |
| *SP670_0702* | 34.42 | 3.07E-50 | 1.86 | 2.63E-07 |
| *SP670_0703* | 24.49 | 1.64E-31 | 1.64 | 1.41E-02 |
| *SP670_0704* | 22.50 | 1.52E-27 | 2.04 | 2.73E-05 |

**Supplemental Table 2.** Primary antibodies used for Microscopy

| Protein | Primary antibody |
|---|---|
| | *Species raised, catalogue and clone, company* |
| JAM-A | α-mouse, SC53623, J10.4, Santa Cruz |
| Claudin 1 | α-mouse, 37-4900, 2H10D10, ThermoFisher |
| Claudin 4 | α-mouse, H00001364, MO2, Biotechne |
| ZO-1 | α-mouse, 339100, Invitrogen |
| β catenin | α-mouse, 2677, L54E2, Cell Signalling Technology |
| Acetylated tubulin | α-mouse, T7451, 6-11B-1, Merck |
| Muc5ac | α-mouse, MAB2011, CLH2, Merck |
| Uterogloblin | α-rabbit, PA5102469, Invitrogen |

**Supplemental Figure 1.** Ciliated Primary HBE-A cells

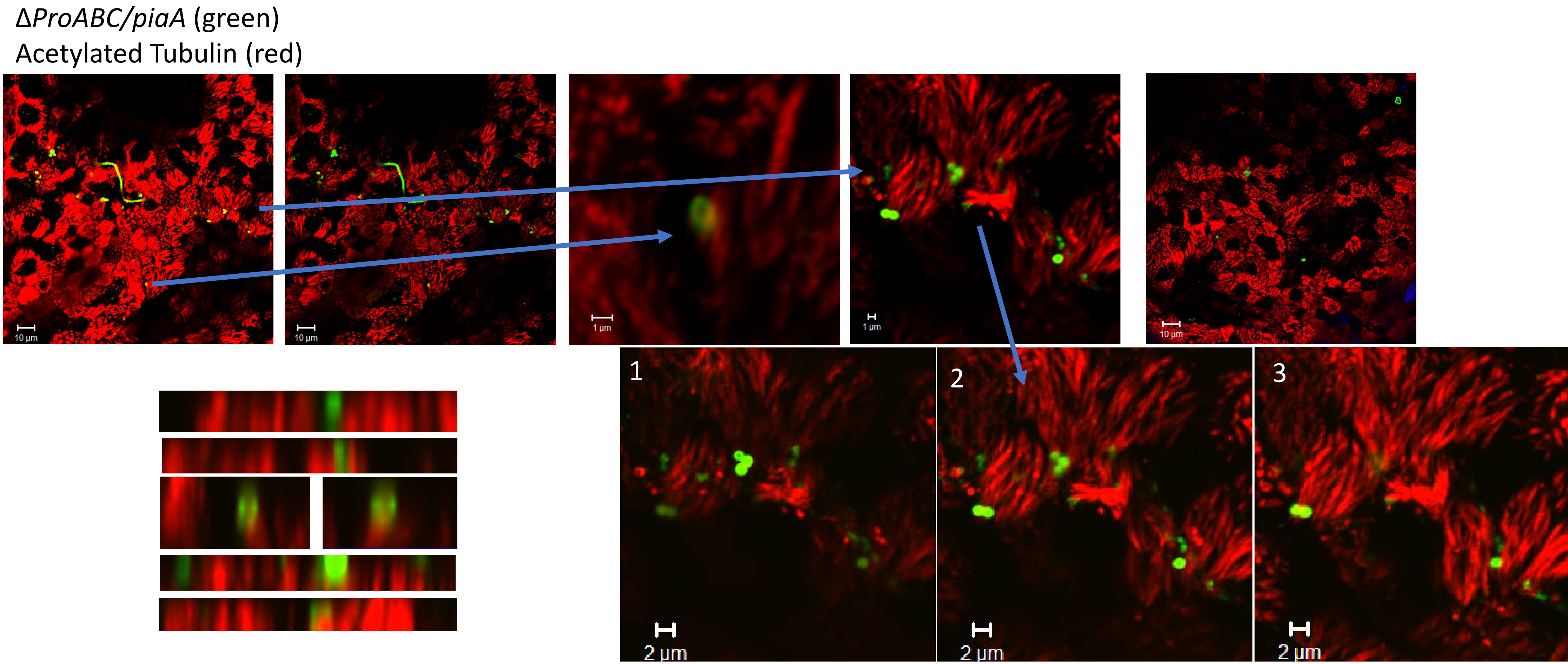

**Supplemental Figure 2.** Association, microinvasion and transmigration of pneumococci following 1hr infection in Detroit 562 cells

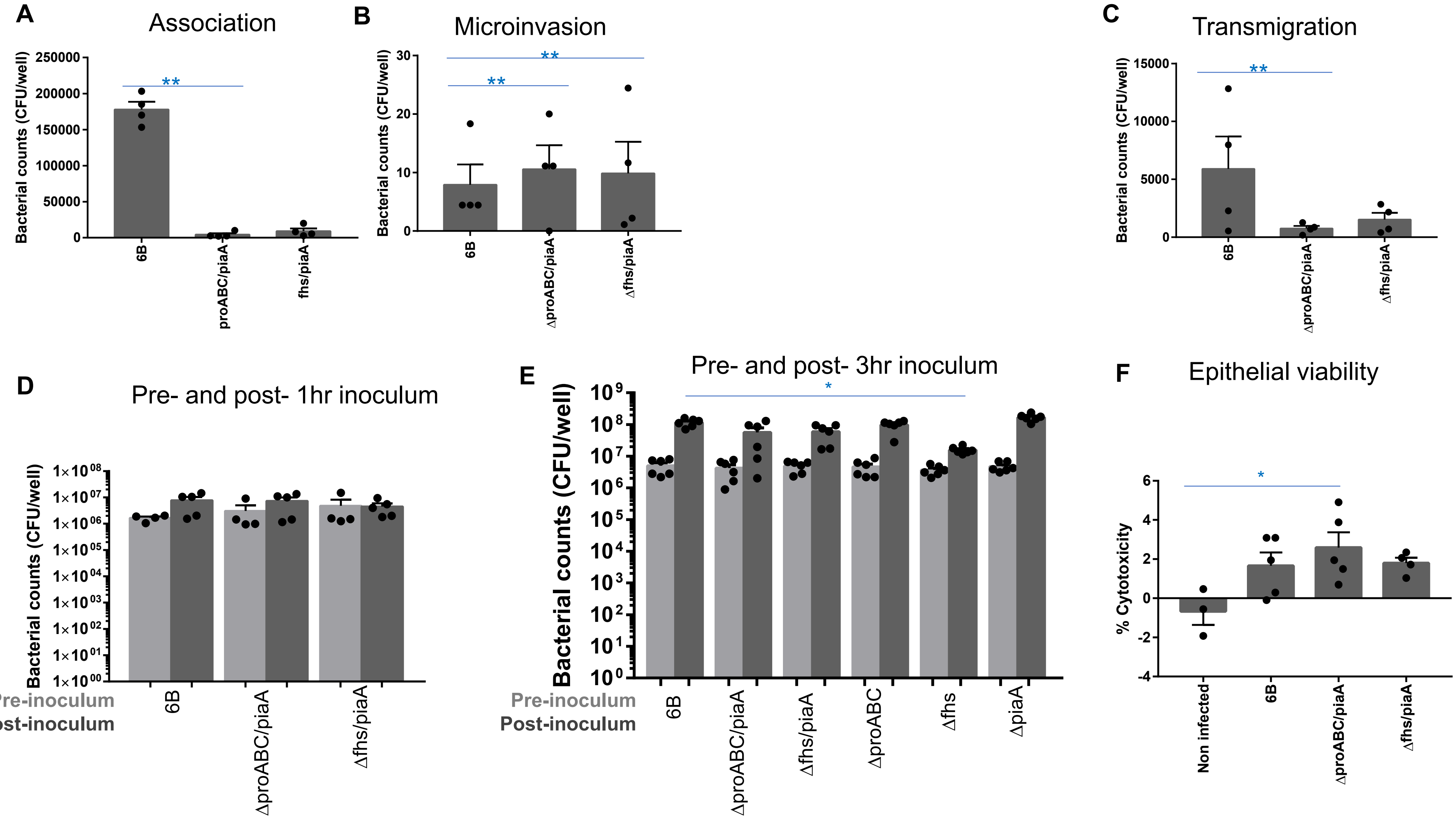
